# Distinct large-scale networks are associated with motor and nonmotor symptoms in Parkinson’s disease

**DOI:** 10.1101/2020.05.13.093724

**Authors:** Sule Tinaz, Serageldin Kamel, Sai S. Aravala, Mine Sezgin, Mohamed Elfil, Rajita Sinha

**Author notes:** Corresponding Author: Sule Tinaz.

## Abstract

Parkinson’s disease (PD) is characterized clinically by various motor and nonmotor symptoms. The underlying neuroanatomical correlates of nonmotor symptoms in PD remain poorly understood. We investigated the differences and commonalities in the neuroanatomical correlates 1) between highly prevalent nonmotor features including fatigue, anxiety, depression, and apathy, and 2) between these nonmotor features and motor severity in nondemented subjects with mild PD (Hoehn & Yahr disease stage 2) using structural and functional magnetic resonance imaging. Compared to matched controls, the PD group showed atrophy in the basal ganglia and superior frontal cortex. Motor severity correlated with cortical thinning in frontotemporal regions, as well as with reduced functional connectivity between the frontostriatal and cerebellar networks. The composite nonmotor symptom severity did not show any correlation with the structural brain data, but correlated with reduced functional connectivity in a large-scale network consisting of frontostriatal, parietotemporal, and cerebellar nodes. The individual components of the nonmotor symptoms also mapped onto specific neural networks. Our study shows that motor and nonmotor features in PD are associated with distinct large-scale networks. The basal ganglia and cerebellum are core regions in all of these networks. The abnormal functional connectivity in the nonmotor network seems to be related to cognitive and emotional dysregulation and may have implications for future cognitive decline in PD.

## INTRODUCTION

The neurodegenerative process in Parkinson’s disease (PD) gradually spreads throughout the whole brain affecting multiple neurotransmitter systems and networks, and causes not only motor, but also a broad spectrum of nonmotor dysfunction (Braak et al., 2003). In fact, several nonmotor features (e.g., hyposmia, constipation, mood disorders, rapid eye movement sleep behavior disorder) can predate the onset of motor symptoms and become increasingly prevalent as the disease progresses (Poewe, 2008). Fatigue and neuropsychiatric disorders including anxiety, depression, and apathy are common nonmotor characteristics of PD and can be present at any stage of the disease diminishing the quality of life of patients (Aarsland et al., 2009, 2011; Weintraub and Burn, 2011; Friedman et al., 2016).

Although fatigue, anxiety, depression, and apathy are separate nonmotor domains, there is considerable overlap between their symptomatology. Significant correlation exists between fatigue and anxiety, apathy, and depression (Skorvanek et al., 2015; Friedman et al., 2016; Siciliano et al., 2018). Apathy and depression share features such as anhedonia, blunted affect, and negative emotions (Pagonabarraga et al., 2015). Depression and comorbid anxiety are also very common (Aarsland et al., 2011).

Molecular imaging studies also suggest commonalities in the affected neurotransmitter systems underlying these nonmotor features in PD. Most, but not all, studies demonstrate deficits or dysregulation in the striatal and limbic dopaminergic, noradrenergic, and serotonergic functions associated with fatigue, anxiety, depression, and apathy in PD (for review, see Yousaf et al., 2017; Valli et al., 2019).

In recent years, there has been an increase in the number of magnetic resonance imaging (MRI) studies investigating network-level structural and functional connectivity changes in PD. In a large cohort of subjects with PD compared to matched controls, a wide network of atrophy including the pedunculopontine nucleus, basal ganglia, thalamus, basal forebrain and limbic structures, and areas in the occipital-temporal-frontal cortices has been demonstrated. Atrophy in these regions also correlated significantly with motor severity (Zeighami et al., 2015). The expression of this atrophy network in PD was found to be more pronounced in a more malignant subtype of PD characterized by higher motor severity and nonmotor symptom scores (Fereshtehnejad et al., 2017).

Functional MRI (fMRI) studies have shown abnormal functional connectivity patterns within and between the nodes of major networks including the sensorimotor, default mode, salience, and executive-attention networks in PD (for a meta-analysis see Tahmasian et al., 2017). A number of MRI studies have also investigated more specifically the neuroanatomical correlates of motor and nonmotor features of PD (for reviews, see Wen et al., 2016; Yousaf et al., 2017; Prell 2018; Valli et al., 2019).

In the motor domain, in general, findings suggest that reduced or abnormal functional connectivity in cortical-striatal-thalamic circuits correlates with motor severity (Cerasa et al., 2016). In addition to total motor severity (Tuovinen et al., 2018), various motor characteristics including tremor (Helmich et al., 2011), freezing of gait (Tessitore et al., 2012), and akinesia (Hensel et al., 2019), as well as motor subtypes of PD such as tremor-dominant and postural instability and gait disorder (Vervoort et al., 2016) have been found to map onto abnormal (increased or decreased) functional connectivity patterns of cortical, cerebellar, and basal ganglia networks.

In the nonmotor domain, fatigue in PD has been found to be associated with atrophy in the dorsal striatum (Kluger et al., 2019), increased functional connectivity between cortical limbic and frontotemporal regions (Zhang et al., 2017) and within the default mode network (Tessitore et al., 2016); and decreased functional connectivity in the sensorimotor network (Tessitore et al., 2016). Apathy in PD has been shown to be associated with atrophy in the nucleus accumbens (Carriere et al., 2014), putamen (Ye et al., 2018) and frontoparietal and limbic regions (Reijnders et al., 2010; Ye et al., 2018), as well as with reduced functional connectivity between predominantly limbic frontostriatal regions (Baggio et al., 2015). Anxiety in PD has been linked to atrophy in the amygdala (Vriend et al., 2016), and anterior cingulate cortex and precuneus (Wee et al., 2016); increased functional connectivity between the left amygdala and parietal regions, and decreased functional connectivity between the left amygdala and frontotemporal regions (Zhang et al., 2019). Depression in PD has been shown to be associated with atrophy in limbic regions including the insula and anterior cingulate (Kostic et al., 2010), and hippocampus and amygdala (van Mierlo et al., 2015). However, increased anterior cingulate volumes have also been found in PD patients with depression (van Mierlo et al., 2015). Correlation between worsening depressive symptoms in PD over time and atrophy in the occipito-temporo-parietal regions and thalamus has been shown (Hanganu et al., 2017). Reports on functional connectivity patterns associated with depression in PD are more variable: Increased functional connectivity between limbic regions (amygdala, mediodorsal thalamus) (Hu et al., 2015), in the posterior cingulate cortex (Lou et al., 2015); in the anterior cingulate cortex, supplementary motor area, and right cerebellum (Wang et al., 2018), between left parahippocampal gyrus and inferior temporal gyrus (Zhang et al., 2019); and decreased functional connectivity between the cortical-limbic networks (Hu et al., 2015); right amygdala and frontoparietal areas (Huang 2015), right middle frontal gyrus and left cerebellum (Wang et al., 2018), and between the left parahippocampal gyrus and orbitofrontal and superior temporal gyri (Zhang et al., 2019) have been observed.

The underlying neuroanatomical correlates of fatigue, anxiety, depression, and apathy in PD remain poorly understood. The heterogeneities in cohort characteristics, subjects’ dopaminergic medication status, and methodological considerations in imaging studies are significant confounding factors. Moreover, in most imaging studies of PD, cognitive and motor status is assessed and reported, but either the presence of subclinical neuropsychiatric/nonmotor symptoms or the considerable overlap between them is not always taken into account in a comprehensive way.

In this study, we aimed to investigate the whole-brain neuroanatomical correlates of motor and nonmotor features (from here on, referring to fatigue, anxiety, depression, and apathy) of the disease in community-dwelling, independent, and non-demented patients with PD using structural and resting-state fMRI. We aimed to address the following question: What are the differences and commonalities in the neuroanatomical mapping 1) between these nonmotor features, and 2) between the motor and nonmotor features? We hypothesized that the nonmotor features will be associated with atrophy and reduced functional connectivity predominantly in the nodes of the limbic cortical-striatal circuits and the motor features will be associated with atrophy and reduced functional connectivity mainly in the nodes of the motor cortical-striatal circuits. We also hypothesized that each nonmotor component will demonstrate a unique neuroanatomical signature.

## METHODS

### Subjects

We included 60 subjects with a diagnosis of idiopathic PD according to the Movement Disorder Society (MDS) criteria (Postuma et al., 2015). All subjects participated in the study after giving written informed consent in accordance with the procedures approved by the Human Research Protection Office of the Yale School of Medicine. Subjects were recruited primarily through the Yale Movement Disorders Clinic and the Connecticut Advocates for Parkinson’s group. The study was conducted at the Yale Magnetic Resonance Research Center. All subjects underwent an initial screening for medical history and MRI safety. We excluded subjects with PD who met any of the following criteria: They were not fully independent (n=1), had a neurological or psychiatric disorder (other than PD and comorbid depression or anxiety), had a medical condition that might affect the central nervous system, had a history of alcohol or illicit drug abuse, had a history of head injury resulting in loss of consciousness, had dementia (Montreal Cognitive Assessment (MoCA) score < 21) (Nasreddine et al., 2005), had contraindications for MRI (n=1), had abnormal findings in anatomical MRI scan (n=1). We used the anatomical MRI scans of 37 age- and gender-matched control (Con) subjects from the Parkinson’s Progression Markers Initiative (PPMI) database. The PPMI is a large, international multicenter clinical study to identify a variety of biomarkers for progression of de novo PD. The database also includes healthy controls who are male or female 30 years or older at screening, with the following exclusion criteria: 1) current or active clinically significant neurological disorder, 2) a first-degree relative with idiopathic PD (parent, sibling, child), 3) a MoCA score of 26 or less; and 4) had received any of the following drugs: neuroleptics, metoclopramide, α-methyldopa, methylphenidate, reserpine, or amphetamine derivative, within six months of screening.

### Clinical data collection and analysis

We assessed disease severity and stage of the PD subjects using the MDS Unified Parkinson’s Disease Rating Scale (MDS-UPDRS) (Goetz et al., 2008) and the Hoehn and Yahr (H & Y) scale (Hoehn and Yahr, 1967). The cutoff for H & Y for inclusion was < 3 (i.e., mild bilateral disease, may have some impairment in balance) to ensure that subjects were fully independent and could tolerate being off medication. We examined the subjects in the morning when they were off dopaminergic medications overnight. Only five subjects were examined in the “medication-on” state, three of whom were not on carbidopa/levodopa treatment. We collected the MRI scans in the morning after subjects took the first dose of their dopaminergic medication.

Following the MDS Task Force recommendations, we administered the Spielberger Trait Anxiety Inventory (STAI-T) (Spielberger et al., 1983, suggested (Leentjens et al., 2008a)), Beck Depression Inventory-II (BDI-II) (Beck et al., 1996, recommended for screening and severity rating (Schrag et al., 2007)), Starkstein Apathy Scale (Starkstein et al., 1992, recommended for screening and severity rating (Leentjens et al., 2008b)), and Parkinson’s Fatigue Scale (PFS-16) (Brown et al., 2005, recommended for screening and severity rating (Friedman et al., 2010)).

### Statistical analysis of the clinical data

We assessed the normality of distribution of the scores using the Shapiro-Wilk test. The mean values and standard deviations of the normally distributed, and median values and median absolute deviations of the non-normally distributed scores were compared with the population mean scores or cutoff scores of the respective tests, when applicable, using one-sample t-tests (p < 0.05, two-tailed). We used the SPSS 26 software for statistical analyses.

### Factor analysis of the clinical data

We first performed pairwise Spearman rho correlation analyses between the STAI-T, BDI-II, apathy, PFS-16, and MDS-UPDRS III scores to examine the degree of collinearity between these variables (see Supplementary Table 3). Given significant collinearity between the STAI-T, BDI-II, apathy, and PFS-16 scores (but not the MDS-UPDRS III scores), we performed dimension reduction using factor analysis in SPSS 26. Principal component analysis with oblique rotation (assumes that the variables are not orthogonal) was used as the extraction method. The cutoff for component extraction was determined based on the eigenvalue > 1. This analysis yielded a single component with the eigenvalue of 2.7 (explaining 67.8% of variance). We called this component the composite nonmotor symptom (NMScomposite) variable.

### MRI data collection

We collected the anatomical MRI scans of 55 and fMRI scans of 43 subjects in a 3.0 Tesla Siemens Trio TIM, and those of five subjects in a 3.0 Tesla Siemens Prisma human research scanner using a 32-channel head coil. The anatomical scans of all Con subjects from the PPMI database were also collected in 3.0 Tesla scanners (29 in Siemens TIM Trio, 4 in Philips Achieva, 3 in General Electric, 1 in Siemens Verio).

We collected high-resolution T1-weighted MPRAGE anatomical images (176 slices, slice thickness: 1 mm, in-plane resolution: 1×1 mm, FoV: 250 mm, Matrix: 256 x 256, TR: 1900 ms, TE: 2.52 ms, TI: 900 ms, flip angle: 9 degrees) for an accurate localization of the fMRI data in the beginning of each scan session. Then, we obtained axial T2-weighted, echo planar functional images at rest for 10 min and 8 s (36 slices, slice thickness: 4 mm, no spacing; in-plane resolution: 3.5×3.5 mm, FoV: 224 mm, Matrix: 64×64, TR: 2000 ms, TE: 25 ms, flip angle: 90 degrees, number of acquisitions: 304). We instructed the subjects to keep their eyes closed, avoid any voluntary movement, and let their mind wander. We assessed wakefulness at the end of the scan by subjects’ report.

### Analysis of the anatomical MRI scans

We used the standard processing pipeline of the FreeSurfer analysis software (available at https://surfer.nmr.mgh.harvard.edu). Briefly, we used the subcortical volume-based stream (Fischl et al., 2002) and the cortical surface-based stream (Dale et al., 1999; Fischl et al, 1999) to obtain the individual subcortical volume and cortical thickness values, respectively. The outcomes of the skull-stripping, cortical tessellation, and subcortical segmentation stages were inspected for quality control using Freeview. Pial surfaces that extended to the dura were edited using the Recon Edit function in Freeview and the surfaces were regenerated subsequently.

### Statistical analysis of the subcortical volumes

We extracted the estimated total intracranial, total brain, total gray matter, cerebral white matter, ventricular, cerebellar, basal ganglia (caudate, putamen, pallidum, accumbens), thalamus, amygdala, and hippocampus volumes. We averaged the basal ganglia, thalamus, amygdala, and hippocampus volumes across the hemispheres. We performed all statistical analyses on volumes that were normalized to the estimated total intracranial volume in SPSS 26. First, we tested a potential age-by-group interaction using a general linear model (GLM) (dependent variable: normalized volume, fixed factor: group with two levels, covariate: age). We then performed between-group comparisons using two-sample t-tests (two-tailed, p < 0.05) when the distribution of the normalized volumes was normal in both Con and PD groups. Levene’s test for equality of variances was also performed. We used nonparametric Mann-Whitney U tests (two-tailed, p < 0.05) when the distribution of the normalized volumes was not normal in either the Con or PD group. We set the statistical significance threshold for seven pairwise comparisons of the subcortical structures at p < 0.007 (0.05/7).

We examined the relationship between the normalized subcortical volumes and the MDS-UPDRS III and NMScomposite scores of the PD group using multiple regression analyses (dependent variable: behavioral score, independent variables: normalized volumes, weighted by age) (two-tailed, p < 0.05).

### Statistical analysis of the cortical surfaces

We assessed the between-group differences in cortical thickness using the Query, Design, Estimate, Contrast (QDEC) module in FreeSurfer. The smoothed cortical surfaces of all subjects from both hemispheres using a Gaussian kernel with full-width half-maximum (FWHM) of 15 mm were the dependent variable and entered into a GLM (categorical variable: group, nuisance variable: age). In separate QDEC analyses, we examined the correlations between cortical thickness and the MDS-UPDRS III and NMScomposite scores of the PD group (nuisance variable: age). For all QDEC analyses, we performed correction for multiple comparisons using Monte Carlo simulations with a significance threshold of 2 corresponding to p < 0.01.

### Analysis of the resting-state fMRI scans

We used the Connectivity toolbox for the resting-state data analysis (Whitfield-Gabrieli and Nieto-Castanon, 2012) of 48 PD subjects that were scanned in medication “on” state. Preprocessing steps included the removal of the first four scans to reach magnetization steady state, motion correction, outlier detection (frame-wise displacement above 0.9 mm or global signal changes above 5 standard deviations), coregistration of functional scans with the anatomical scan, normalization to the standard Montreal Neurological Institute (MNI) brain template, and smoothing with a Gaussian kernel with a FWHM of 8 mm to account for interindividual anatomical variability. De-noising steps included the elimination of signal originating from the white matter and cerebrospinal fluid, regression of motion artifacts and outliers from the time series, scrubbing, and quadratic detrending. Global signal was not removed. Finally, we bandpass-filtered (0.008 < f < 0.1 Hz) the data to capture the fluctuations of the blood oxygenation level-dependent (BOLD) signal that typically occur within this frequency range at rest.

We used the functionally defined nodes (n=268) of the whole-brain Shen atlas (Shen et al., 2013) for the functional connectivity analyses. For each subject, we extracted the average BOLD signal time courses from these nodes and correlated them with each other using Pearson correlations. The “r” values corresponded to the functional connectivity strength between node pairs. We Fisher z-transformed the “r” values and obtained group-level functional connectivity maps for statistical analyses. Finally, we used the MDS-UPDRS III and NMScomposite scores as the covariates of interest and age as a covariate of no interest in separate models to be correlated with these maps. In exploratory analyses, we also used the components of the NMScomposite (i.e., STAI-T, BDI-II, apathy, PFS-16 scores) separately as covariates of interest and age as a covariate of no interest for correlation with these maps. We used the false discovery rate (FDR) method for correction for multiple comparisons (p < 0.05) (Genovese et al., 2002).

## RESULTS

### Demographic and clinical data

The mean age of the Con group (n = 37) was 65.8 ± 8.7 years (range: 45.6 – 82) (Shapiro-Wilk test p = 0.939) and that of the PD group (n = 60) was 64.6 ± 9.1 years (range: 44 – 79.7) (Shapiro-Wilk test p = 0.181). There were 23 males and 14 females in the Con group and 36 males and 24 females in the PD group. The groups did not differ significantly in age (two-sample t-test, p = 0.529, equal variances) or gender (Mann-Whitney U test, p = 0.833). The H & Y score was 2 in all but one (2.5) PD subjects (2.01 ± 0.06). The symptom onset side was left in 28 and right in 32 PD subjects. Ten PD subjects were left-handed. Four PD subjects were not on any medication, 39 were on carbidopa/levodopa, 20 were on dopamine receptor agonists, 34 were on MAO-B inhibitors, 15 were on amantadine, one was on trihexyphenidyl, and one was on Mucuna powder. Nineteen PD subjects were only on one medication. Three subjects were on a combination of four, 11 subjects on three, and 23 subjects on two medications. Time between last dose of dopaminergic medication and motor exam was 14.4 ± 3.8 hr. Of note, 5 subjects were examined in “on” state (1.8 ± 0.8 hr), 3 of whom were not on carbidopa/levodopa. The clinical data of the PD group are summarized in Table 1 (See also Supplementary Material Table 1 for the PD subgroup of n = 48). The details of the behavioral analysis results are summarized in Supplementary Material Table 2. Briefly, our PD cohort scored significantly below the apathy (14), minimal depression (13), and fatigue (2.95) cutoff scores; and significantly above the MoCA cutoff score (26). The trait-anxiety scores of the females in the PD group were also significantly below, whereas those of the males were slightly above (trend) the age- and gender-matched population means. In the PD subgroup (n = 48), however, the trait-anxiety of males was significant.

**Table 1.** Clinical data of the PD group (N=60)

| | Mean ± SD (range) | S-W test p values |
| --- | --- | --- |
| Age (yr) | $64.6 \pm 9.1$ (44 – 79.7) | 0.181 |
| Disease duration (yr) | $5.6 \pm 3.7$ (0.21 – 14.60) | 0.019 |
| LEDD (mg) | $497.6 \pm 313.1$ (100-1640) | 0.000 |
| MDS-UPDRS I | $7.6 \pm 4.4$ (0 – 19) | 0.161 |
| MDS-UPDRS II | $9.8 \pm 5.2$ (1 – 23) | 0.087 |
| MDS-UPDRS III (n = 60) | $30.1 \pm 8.5$ (12 – 58) | 0.299 |
| MDS-UPDRS III off (n = 55) | $30.1 \pm 8.7$ (12 – 58) | 0.287 |
| MDS-UPDRS III on (n = 5) | $30.2 \pm 6.8$ (21 – 36) | 0.142 |
| MDS-UPDRS IV | $1.8 \pm 2.2$ (0 – 13) | 0.000 |
| MDS-UPDRS total | $49.3 \pm 12.8$ (25 – 83) | 0.119 |
| MoCA | $27.5 \pm 2.0$ (23 – 30) | 0.000 |
| STAI-T | $34.1 \pm 10.2$ (21 – 63) | 0.000 |
| BDI-II | $7.7 \pm 6.9$ (0 – 36) | 0.000 |
| Apathy | $8.8 \pm 5.6$ (0 – 22) | 0.007 |
| PFS-16 | $2.3 \pm 0.9$ (0.5 – 4.8) | 0.140 |

BDI-II: Beck depression inventory-II, LEDD: Levodopa equivalent daily dose, MDS-UPDRS: Movement Disorders Society-Unified Parkinson's Disease Rating Scale (part III: Motor exam, on: on medication, off: off medication), MoCA: Montreal cognitive assessment test, PFS-16: Parkinson's fatigue scale (averaged scores), STAI-T: Spielberger State and Trait Anxiety Inventory – Trait, S-W: Shapiro-Wilk test for normality of distribution.

### Subcortical volumetric results

Table 2 shows the results of the Shapiro-Wilk tests for normality of distribution, Mann-Whitney U nonparametric tests for comparison of volumes that were not normally distributed in either group, and two-sample t-tests for comparison of volumes that were normally distributed in both groups. There was no significant age-by-group interaction in any of the volumes. The PD group demonstrated significantly lower volume in the putamen, amygdala, and nucleus accumbens, and significantly higher volume in the pallidum compared to the Con group. The raw subcortical volumes are listed in the Supplementary Material Table 4.

**Table 2.** Subcortical volumetric results

| Volume (mm<sup>3</sup>) | Con (N = 37) | PD (N = 60) | S-W p values |  | M-W/t-test |
| --- | --- | --- | --- | --- | --- |
|  |  |  | Con | PD | p values |
| Total intracranial | 1,519,807 ± 170,334 | 1,588,293 ± 147,266 | 0.163 | 0.980 | 0.039 |
| Total brain | 1,126,181 ± 108,658 | 1,188,427 ± 107,975 | 0.000 | 0.468 | 0.073 |
| GM | 612,695 ± 55,390 | 630,768 ± 55,336 | 0.007 | 0.127 | 0.444 |
| WM | 449,229 ± 66,558 | 440,504 ± 150,971 | 0.000 | 0.000 | 0.218 |
| WM hypointensity | 3,494 ± 5,010 | 2,287 ± 1,980 | 0.000 | 0.000 | 0.145 |
| Cerebellum (GM) | 105,419 ± 11,636 | 112,439 ± 11,976 | 0.190 | 0.201 | 0.427 |
| Thalamus | 6,851 ± 775 | 7,398 ± 800 | 0.109 | 0.655 | 0.098 |
| Caudate | 3,425 ± 396 | 3,415 ± 415 | 0.053 | 0.071 | 0.041 |
| Putamen | 4,598 ± 631 | 4,482 ± 570 | 0.258 | 0.378 | 0.006* |
| Pallidum | 1,964 ± 205 | 2,275 ± 318 | 0.010 | 0.000 | 0.000* |
| Hippocampus | 4,004 ± 350 | 4,013 ± 375 | 0.004 | 0.309 | 0.220 |
| Amygdala | 1,669 ± 202 | 1,583 ± 210 | 0.038 | 0.719 | 0.002* |
| Accumbens | 469 ± 87 | 388 ± 77 | 0.069 | 0.461 | 0.000* |

GM: Gray matter, M-W: Mann-Whitney U nonparametric test for volumes that were not normally distributed in either group, S-W: Shapiro-Wilk test for normality of distribution, WM: White matter. All statistical tests were performed on volumes that were averaged and normalized to the total intracranial volume. \*: Significant p values that are below the statistical threshold of $p < 0.007$ (0.05/7).

### Correlations between the behavioral and subcortical volumes

The multiple regression analyses did not reveal any significant correlation between the normalized subcortical volumes and the MDS-UPDRS III or the NMScomposite scores. See the scatterplots in Supplementary Material Figure 1 for details.

### Cortical thickness results

There was no significant age-by-group interaction in cortical thickness of any region. PD group showed significant cortical atrophy in frontal regions bilaterally compared to the Con group (Figure 1, Table 3). There was also a significant negative correlation between the MDS-UPDRS III scores and right temporal and frontal cortical thickness in the PD group (Figure 2, Table 3). There was no significant correlation between cortical thickness in any area and the NMScomposite scores or any of its individual components.

**Figure 1.**
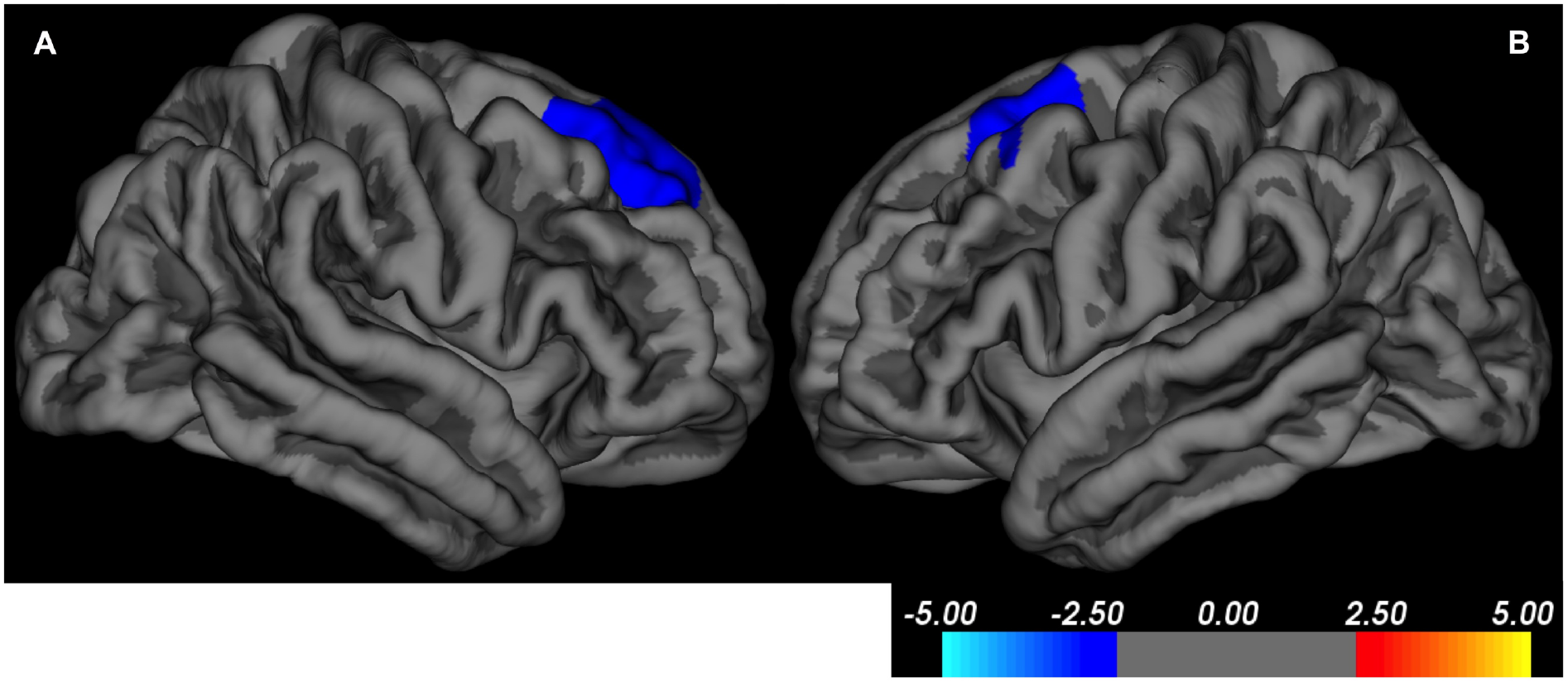
Cortical thinning in the PD compared to the Con group in **A**: Right superior frontal cortex (BA8) and **B**: Left premotor cortex (BA6). Color bar displays the t values (p < 0.01, corrected for multiple comparisons using Monte-Carlo simulations).

**Figure 2.**
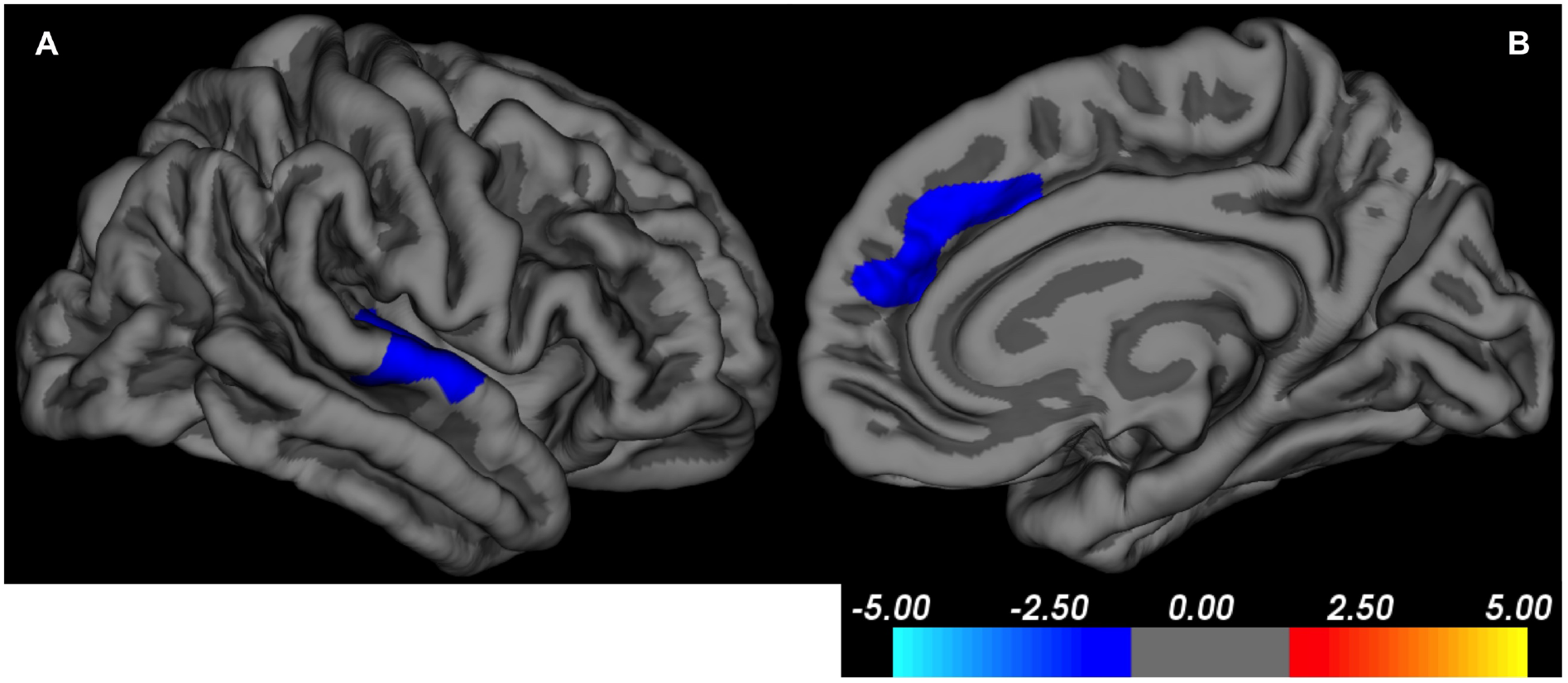
The MDS-UPDRS III scores in the PD group correlate with cortical thinning in **A**: Right superior temporal cortex (BA22) and **B**: Right medial superior frontal cortex (BA8). Color bar displays the t values (p < 0.01, corrected for multiple comparisons using Monte-Carlo simulations).

**Table 3.** Cortical thickness results

|  | MNI coordinates |  |  | BA | Cluster size (mm<sup>2</sup>) | T |
| --- | --- | --- | --- | --- | --- | --- |
|  | x | y | z |  |  |  |
| **PD < Con** |  |  |  |  |  |  |
| Left premotor | -19 | 13 | 57 | 6 | 1071 | -4.05 |
| Right superior frontal | 19 | 41 | 35 | 9 | 1115 | -3.15 |
| **Correlation with MDS-UPDRS III** |  |  |  |  |  |  |
| Right superior temporal | 67 | -14 | -1 | 22 | 842 | -2.98 |
| Right superior frontal | 10 | 22 | 37 | 8 | 739 | -3.83 |

BA: Brodmann area, Con: Control, MNI: Montreal Neurological Institute, PD: Parkinson's disease ( $p < 0.01$ , corrected for multiple comparisons using Monte-Carlo simulations).

### Resting-state fMRI results

The time between last dose of dopaminergic medication and start of resting-state fMRI scan was 185 ± 68 min (range: 40-315 min).

Head motion averaged across x, y, z planes was 0.29 ± 0.36 mm. Head rotation averaged across x, y, z planes was 0.28 ± 0.38 degrees. There were on average 1.9 ± 5.3 invalid scans (see Supplementary Material Table 5 for details). These results show that our PD cohort had minimal head motion.

The MDS-UPDRS III scores correlated negatively with the pairwise functional connectivity predominantly between the caudal and middle parts of the right caudate and prefrontal cortical regions and cerebellum. The functional connectivity between the right temporal pole and prefrontal cortical regions correlated positively with the MDS-UPDRS III scores (Figure 3, Table 4). The NMScomposite scores correlated negatively with the pairwise functional connectivity predominantly between the basal ganglia and frontotemporal regions, dorsal posterior cingulate cortex, and cerebellum. The functional connectivity of the basal ganglia and cerebellum with the occipitotemporal regions correlated positively with the NMScomposite scores (Figure 4, Table 5).

**Figure 3.**
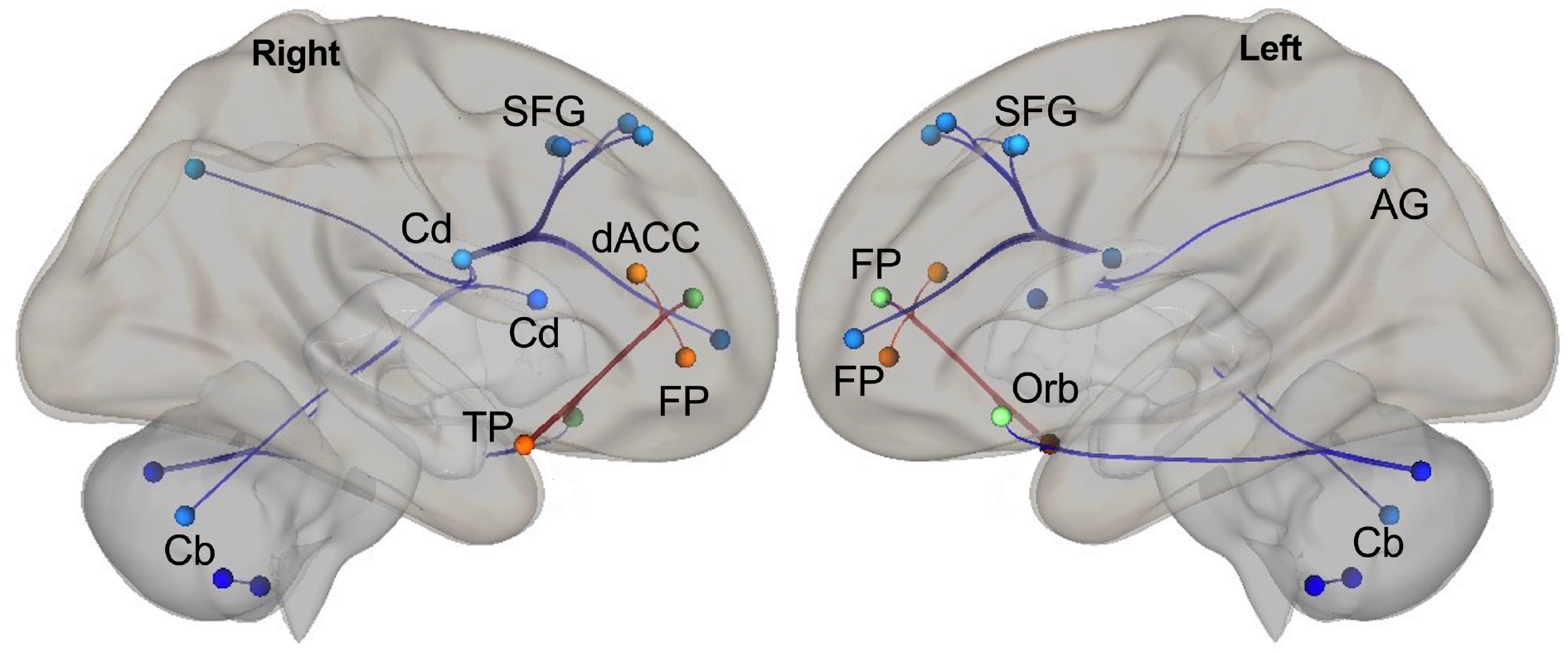
Correlations between the MDS-UPDRS III scores and pairwise functional connectivity. (FDR-corrected p < 0.05, warm colors: positive, cool colors: negative). Nodes in both hemispheres are displayed from right- and left-view on the MNI template. AG: Angular gyrus, Cb: Cerebellum, Cd: Caudate, dACC: Dorsal anterior cingulate cortex, FP: Frontal pole (anterior prefrontal cortex), Orb: Orbitofrontal cortex, SFG: Superior frontal gyrus (medial and lateral), TP: Temporal pole.

**Table 4.** Correlations between MDS-UPDRS III scores and pairwise functional connectivity

| *Negative Correlations* |  |  |  |  |
| --- | --- | --- | --- | --- |
| *Node Pairs* |  | *Pair Labels* |  |  |
| Numbers | Node 1 (BA) | Node 2 (BA) | T | p |
| (241)-(186) | L cerebellum | L TP (BA38) | -4.54 | 0.011 |
| (122)-(142) | R caudate | L aPFC (BA10) | -4.39 | 0.018 |
| (122)-(140) | R caudate | L aPFC (BA10) | -4.15 | 0.019 |
| (122)-(12) | R caudate | R SFG (BA8) | -3.98 | 0.019 |
| (122)-(182) | R caudate | L AG (BA39) | -3.94 | 0.019 |
| (109)-(249) | R cerebellum | R cerebellum | -4.27 | 0.027 |
| (122)-(148) | R caudate | L SFG (BA8) | -3.67 | 0.033 |
| (122)-(105) | R caudate | R cerebellum | -3.62 | 0.033 |
| (121)-(241) | R caudate | L cerebellum | -3.92 | 0.040 |
| (122)-(150) | R caudate | L SFG (BA8) | -3.50 | 0.040 |
| (122)-(149) | R caudate | L SFG (BA8) | -3.43 | 0.044 |
| (241)-(153) | L cerebellum | L IFG (BA47) | -3.75 | 0.045 |

**Positive Correlations**
| *Node Pairs* | *Pair Labels* |  |  |  |
| --- | --- | --- | --- | --- |
| Numbers | Node 1 (BA) | Node 2 (BA) | T | p |
| (53)-(140) | R TP (BA38) | L aPFC (BA10) | 4.54 | 0.011 |
| (53)-(153) | R TP (BA38) | L IFG (BA47) | 3.81 | 0.041 |
| (53)-(83) | R TP (BA38) | R dACC (BA32) | 3.78 | 0.041 |
| (53)-(5) | R TP (BA38) | R aPFC (BA10) | 3.64 | 0.046 |

AG: Angular gyrus, aPFC: Anterior prefrontal cortex, BA: Brodmann area, dACC: Dorsal anterior cingulate cortex, IFG: Inferior frontal gyrus, orbital part, SFG: Superior frontal gyrus, TP: Temporal pole. See the interactive webpage:
https://bioimagesuiteweb.github.io/webapp/connviewer.html for the coordinates of the Shen Atlas nodes (Shen et al., 2013).

**Figure 4.**
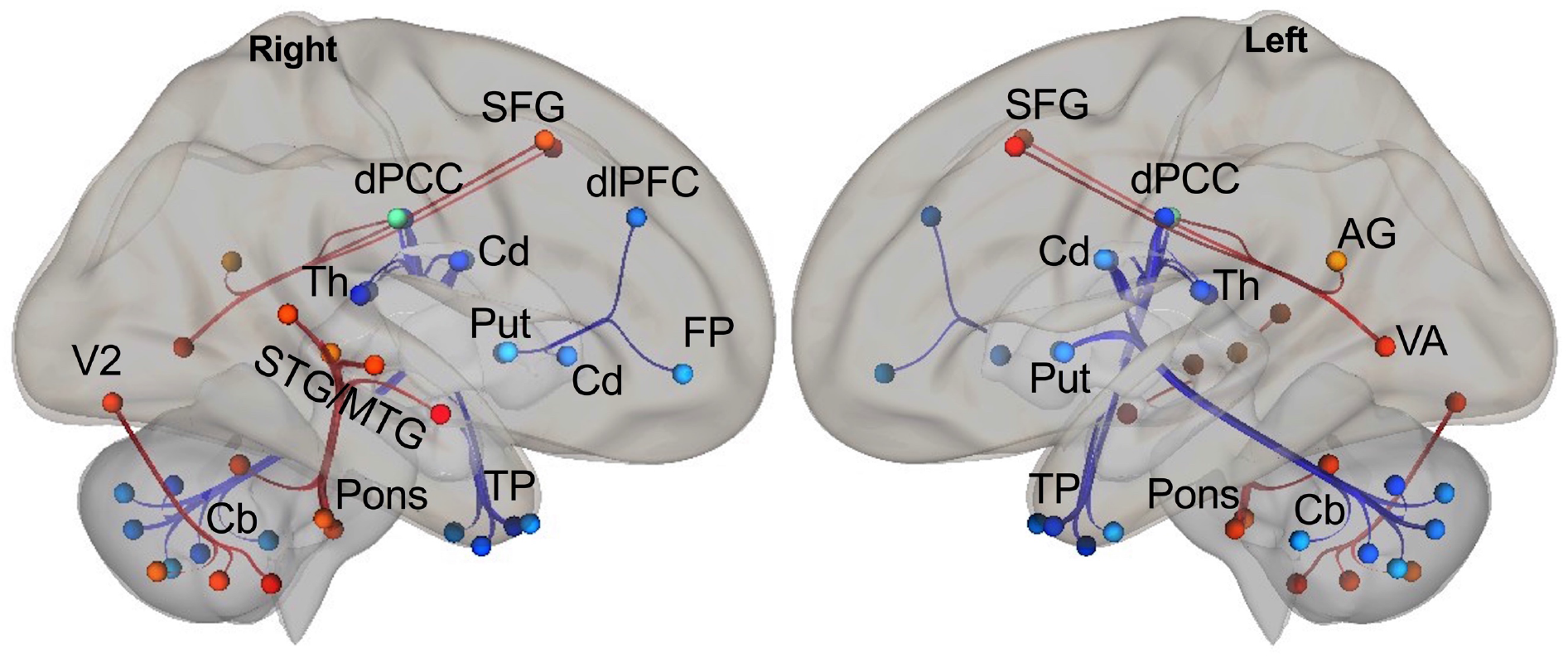
Correlations between the NMScomposite scores and pairwise functional connectivity. (FDR-corrected p < 0.05, warm colors: positive, cool colors: negative). Nodes in both hemispheres are displayed from right- and left-view on the MNI template. AG: Angular gyrus, Cb: Cerebellum, Cd: Caudate, dlPFC: Dorsolateral prefrontal cortex, dPCC: Dorsal posterior cingulate cortex, FP: Frontal pole (anterior prefrontal cortex), MTG: Middle temporal gyrus, Put: Putamen, SFG: Superior frontal gyrus (medial and lateral), STG: Superior temporal gyrus, TP: Temporal pole, Th: Thalamus, V2: Secondary visual area, VA: Visual association cortex.

**Table 5.**
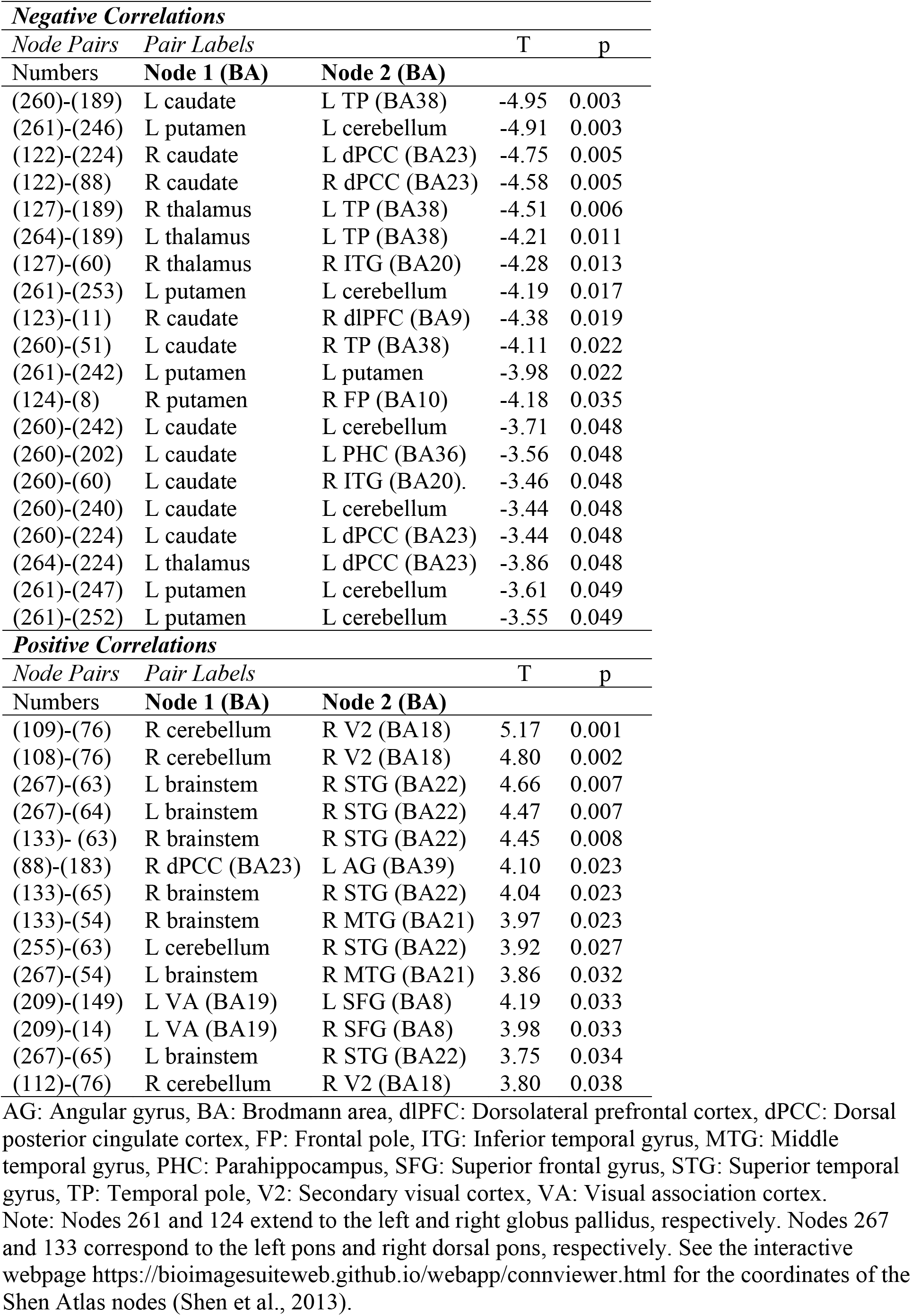
Correlations between NMScomposite scores and pairwise functional connectivity

The individual components of the nonmotor symptoms showed distinct correlation patterns with the pairwise functional connectivity across the whole brain. Apathy scores correlated negatively with the functional connectivity of the basal ganglia with the prefrontal regions and cerebellum, as well as of the fusiform gyrus with numerous other, mainly limbic temporal regions (e.g., insula, hippocampus, parahippocampus, amygdala) (see Supplementary Material Figure 2 and Table 6). Depression scores correlated positively with the functional connectivity of the anterior cingulate cortex and insula with sensorimotor regions, as well as with the functional connectivity between frontoparietal regions. Depression scores showed relatively few negative correlations with the functional connectivity between the basal ganglia and limbic frontotemporal regions (see Supplementary Material Figure 3 and Table 7). Fatigue scores correlated positively with the functional connectivity between the occipitotemporal regions and brainstem and cerebellum (to a lesser extent also the basal ganglia). Relatively few negative correlations were observed between the fatigue scores and functional connectivity of the basal ganglia with the cerebellum and temporal regions (see Supplementary Material Figure 4 and Table 8). Anxiety scores showed a small number of correlations, negative with the functional connectivity between the putamen and cerebellum, and positive with the functional connectivity between the cerebellum and occipital regions (see Supplementary Material Figure 5 and Table 9).

## DISCUSSION

We demonstrated significant collinearity between the nonmotor features including fatigue, apathy, depression, and anxiety in our PD group. Our factor analysis justified the grouping of these features as a composite nonmotor variable. Of note, none of the nonmotor features showed any significant correlation with motor severity, therefore, significant interaction between the motor and nonmotor domains was not a concern in the separate correlation analyses with the imaging data.

We found atrophy in the prefrontal cortex, putamen, nucleus accumbens, and amygdala in our cohort of nondemented subjects with mild bilateral PD compared to matched controls. Motor severity correlated with focal atrophy in the right frontotemporal cortex, but not with any of the subcortical volumes. These findings are in line with previous reports of cortical and subcortical atrophy pattern in PD and its relationship to motor severity (Zeighami et al., 2015; Fereshtehnejad et al., 2017). Nonmotor symptoms of PD have been linked to atrophy in the frontostriatal regions and limbic cortical and subcortical structures (Kostic et al., 2010; Reijnders et al., 2010; Carriere et al., 2014; van Mierlo et al., 2015; Vriend et al., 2016; Wee et al., 2016; Hanganu et al., 2017; Ye et al., 2018; Kluger et al., 2019). However, we found no significant correlation between the composite or individual nonmotor symptom scores and structural brain data. A possible explanation for this discrepancy is that, unlike in most previous studies, our PD cohort as a whole did not have significant fatigue, apathy, depression, or anxiety. Moreover, we used a whole-brain approach, but not all previous reports were based on whole-brain structural data and focused on a limited number of regions selected a priori (Carriere et al., 2014; van Mierlo et al., 2015; Vriend et al., 2016; Wee et al., 2016; Kluger et al., 2019).

Consistent with our hypothesis, the motor and nonmotor features of our PD cohort mapped onto distinct functional brain networks. Motor severity correlated predominantly with reduced functional connectivity not only in the motor, but also in the cognitive frontostriatal and cerebellar circuits. Frontal nodes of these circuits, especially in the superior frontal cortex, also overlapped with areas of significant atrophy. Though we did not specifically predict it, the cerebellar involvement is not surprising given its rich connections with the basal ganglia and frontal cortex (Bostan et al., 2013). Cerebellar hyperactivity and hyperconnectivity have been reported in task-based and resting-state fMRI studies of PD (Helmich et al., 2011, Wu and Hallett, 2013, Festini et al., 2015) and generally attributed to compensatory mechanisms. This functional pattern was most consistently exhibited in PD subjects scanned off dopaminergic medication and was either normalized or even reduced by medication (Festini et al., 2015; Solstrand Dahlberg et al., 2020). Moreover, different cerebellar lobules have been found to display diverse activation and connectivity characteristics in PD depending on the motor, emotional, and cognitive context (Solstrand Dahlberg et al., 2020) as we have also observed in our motor and nonmotor network data.

The severity of the nonmotor feature cluster correlated with reduced functional connectivity in a large-scale network consisting of frontostriatal, parietotemporal, and cerebellar nodes. Two nodes specifically warrant further discussion: The temporal pole comprises subregions with distinct functional connectivity profiles. Our temporal pole node overlaps with the anterior areas 35/36 in Pascual et al. (2015), which exhibit resting-state functional connectivity with paralimbic cortical, limbic subcortical, and occipitotemporal regions, and are thought to mediate higher-order visual and semantic processing, assessment of the value and relevance of visual information, and eye movement control (Pascual et al., 2015). The posterior cingulate cortex node in our study overlaps with the dorsal portion of BA23. In addition to being a hub in the default mode network involved in internally-oriented mental processes, the dorsal portion is also strongly connected to frontoparietal task-positive networks involved in cognitive control, and is thought to dynamically regulate the focus of attention (Leech and Sharp, 2014). Our results suggest that dysfunction in a large-scale network that mediates cognitive and limbic functions underlies the nonmotor symptoms in PD. Importantly, two hubs of this network, namely the temporal pole and posterior cingulate cortex, are affected in semantic dementia (Acosta-Cabronero et al., 2011) and Alzheimer’s disease (Buckner et al., 2008), respectively, implying that the dysregulation of this network may also be a harbinger of cognitive decline. This is supported by a parietotemporal pattern of atrophy that has been associated with worse cognitive performance in nondemented PD subjects (Uribe et al., 2016; 2018), as well as correlation between cognitive impairment in PD and reduced functional connectivity in the default mode network, and frontoparietal and temporal regions (Wolters et al., 2019).

Despite being strongly related at the behavioral level, the individual components of the nonmotor features demonstrated distinct network correlates. Among all components, anxiety exhibited the weakest functional connectivity pattern, which was essentially indistinguishable from the others. The reduced functional connectivity in the frontostriatal and cerebellar networks associated with apathy seems to have contributed to the composite nonmotor pattern. Interestingly, apathy also correlated significantly with decreased functional connectivity between the fusiform gyrus, which is involved in facial recognition, and limbic medial temporal regions. This finding is concordant with reports of impaired facial emotion recognition (Enrici et al., 2015; Kalampokini et al., 2018) and its relationship to apathy in nondemented subjects with PD (Martínez-Corral et al., 2010; Robert et al., 2014).

In contrast to apathy, decreased functional connectivity associated with depression and fatigue was relatively rare, however, consistently involved the basal ganglia, cerebellum, and frontotemporal regions. Depression correlated with increased functional connectivity between the limbic anterior cingulate cortex/insula and sensorimotor networks. Increased functional connectivity (Wang et al., 2018) and increased metabolic activity in similar regions (Wen et al., 2016) associated with depression in PD have been reported.

The brainstem, midbrain, and cerebellar functional connectivity with occipitotemporal regions correlated significantly with fatigue and seems to have contributed to the composite nonmotor pattern that exhibited increased functional connectivity in similar regions. The reticular formation is a set of interconnected neurons and spans the brainstem and midbrain. It includes ascending pathways to the thalamus and cortex, projections to the cerebellum, and descending pathways to the spinal cord. A major function of this system is to modulate arousal and consciousness (Zeman, 2001). Our results suggest that the “overdrive” in this network may be associated with fatigue in PD. Alternatively, it is also possible that this “overdrive” actually reflects the effort of subjects to stay awake during the resting-state scan.

In conclusion, our study shows that motor and nonmotor features are linked to distinct large-scale networks in a cohort of nondemented subjects with mild PD and subclinical nonmotor symptoms. The basal ganglia and cerebellum are core regions in all of these networks. While the individual nonmotor features have their unique neural signature, they also converge on a network composed of frontostriatal, parietotemporal, and cerebellar nodes. The abnormal functional connectivity within this network seems to be related to cognitive and emotional dysregulation and may have implications for future cognitive decline in PD.

### Limitations

We scanned our subjects in “on” medication state, but not necessarily at the peak of the dopaminergic medication dose. This may have introduced some variability to the functional connectivity data. In addition, our results should be interpreted with caution and not be generalized as the “off” medication functional connectivity would be expected to show considerable differences. Due to the lack of separate control groups who do not have PD, but exhibit the nonmotor symptoms, we cannot disentangle the contributions of PD-related pathological processes from those underlying the specific nonmotor symptoms to our network findings. Therefore, we can only claim correlations with regard to the degree of these symptoms.

## Acknowledgment

We thank the Connecticut Advocates for Parkinson’s advocacy group for their help with recruitment. Structural MRI data used in the preparation of this article were obtained from the Parkinson’s Progression Markers Initiative (PPMI) database (www.ppmi-info.org/data). For up-to-date information on the study, visit www.ppmi-info.org/data. The PPMI, a public-private partnership, is funded by the Michael J. Fox Foundation for Parkinson’s Research and funding partners, including Abbvie, Avid, Biogen, Bristol-Myers Squibb, Covance, GE Healthcare, Genentech, GlaxoSmithKline, Lilly, Lundbeck, Merck, Meso Scale Discovery, Pfizer, Piramal, Roche, Sanfo Genzyme, Servier, Takeda, Teva, UCB, and Golub Capital.

## Funding

This work was supported by the K23 NS099478 grant from the National Institute of Neurological Disorders and Stroke.

## Notes

### Competing Interest Statement

The authors have declared no competing interest.

