## Supplemental Material for "Distinct large-scale networks are associated with motor and nonmotor symptoms in Parkinson’s disease"

**Supplementary Material**

Forty-eight out of 60 PD subjects underwent resting-state fMRI scanning in medication "on" state. The H & Y score was 2 in all subjects. The symptom onset side was left in 21 and right in 27 subjects. Four subjects were left-handed. Four PD subjects were not on any medication, 31 were on carbidopa/levodopa, 14 were on dopamine receptor agonists, 27 were on MAO-B inhibitors, 12 were on amantadine, and one was on trihexyphenidyl. Fourteen subjects were only on one medication. One subject was on a combination of four, ten subjects on three, and 17 subjects on two medications. Table 1 summarizes the clinical data of this subgroup separately. Time between last dose of dopaminergic medication and motor exam was 15.0 ± 3.8 hr. Of note, 5 subjects were examined in "on" state (1.8 ± 0.8 hr), 3 of whom were not on carbidopa/levodopa.

**Table 1. Clinical data of the PD group (N=48)**

| | Mean ± SD (range) | S-W test p |
|---|---|---|
| Age (yr) | 65.2 ± 8.6 (45.3 – 79.7) | 0.470 |
| Disease duration (yr) | 5.6 ± 3.9 (0.2 – 14.6) | 0.024 |
| LEDD (mg) | 505.2 ± 332.4 (100 – 1640) | 0.000 |
| MDS-UPDRS I | 8.0 ± 4.3 (0 – 19) | 0.530 |
| MDS-UPDRS II | 9.6 ± 5.4 (1 – 23) | 0.097 |
| MDS-UPDRS III (N = 48) | 29.4 ± 8.4 (12 – 58) | 0.117 |
| MDS-UPDRS III off (N = 43) | 29.4 ± 8.7 (12 – 58) | 0.102 |
| MDS-UPDRS III on (N = 5) | 30.2 ± 6.8 (21 – 36) | 0.162 |
| MDS-UPDRS IV | 1.8 ± 2.3 (0 – 13) | 0.000 |
| MDS-UPDRS total | 48.9 ± 12.8 (25 – 83) | 0.088 |
| MoCA | 27.7 ± 1.9 (23 – 30) | 0.000 |
| STAI-T | 34.5 ± 10.3 (21 – 63) | 0.000 |
| BDI-II | 7.4 ± 6.7 (0 – 36) | 0.000 |
| Apathy | 8.8 ± 5.2 (1 – 20) | 0.007 |
| PFS-16 | 2.3 ± 0.9 (1 – 4) | 0.028 |

BDI-II: Beck depression inventory-II, LEDD: Levodopa equivalent daily dose, MDS-UPDRS: Movement Disorders Society-Unified Parkinson's Disease Rating Scale (part III: Motor exam, on: on medication, off: off medication), MoCA: Montreal cognitive assessment test, PFS-16: Parkinson's fatigue scale (averaged scores), STAI-T: Spielberger State and Trait Anxiety Inventory – Trait, S-W: Shapiro-Wilk test for normality of distribution.

**Statistical analyses and results of behavioral data**

The mean values and standard deviations (SD) of the normally distributed (Shapiro-Wilk $p > 0.05$), and median values and median absolute deviations (MAD) of the non-normally distributed (Shapiro-Wilk $p < 0.05$) scores were compared with population mean or cutoff scores of the respective tests using one-sample t-tests ($p < 0.05$, two-tailed). Table 2a (n=60) and 2b (n=48) show the results.

**Table 2a. Behavioral statistics (N=60)**

| | Shapiro-Wilk *p values* | Mean ± SD or Median ± MAD | Population mean or cutoff scores | one-sample t-test *p values* |
|---|---|---|---|---|
| Apathy | 0.007 | 7.0 ± 4.0 | 14.0 | 0.000 |
| STAI-T | 0.011 (M)<br>0.000 (F) | 35.5 ± 6.5 (M),<br>28.5 ± 3.5 (F) | 33.86 ± 8.86 (M),<br>31.79 ± 7.78 (F) | 0.055 (M),<br>0.000 (F) |
| PFS-16 | 0.140 | 2.3 ± 0.9 | 2.95 | 0.000 |
| BDI-II | 0.000 | 6.0 ± 4.0 | 13.0 (minimal depression) | 0.000 |
| MoCA | 0.000 | 28.0 ± 1.0 | 26 | 0.000 |

**Table 2b. Behavioral statistics (N=48)**

| | Shapiro-Wilk *p values* | Mean ± SD or Median ± MAD | Population mean or cut-off scores | one-sample t-test *p values* |
|---|---|---|---|---|
| Apathy | 0.007 | 7.0 ± 4.0 | 14.0 | 0.000 |
| STAI-T | 0.007 (M)<br>0.006 (F) | 37.0 ± 8.0 (M),<br>29 ± 3 (F) | 33.86 ± 8.86 (M),<br>31.79 ± 7.78 (F) | 0.009 (M),<br>0.000 (F) |
| PFS-16 | 0.028 | 2.3 ± 0.7 | 2.95 | 0.000 |
| BDI-II | 0.000 | 6.0 ± 3.0 | 13.0 (minimal depression) | 0.000 |
| MoCA | 0.000 | 28.0 ± 1.0 | 26 | 0.000 |

**Distribution of Behavioral Data**

Out of 60 subjects with PD:

- Eleven had a MoCA score < 26. None had a score < 21.
- Twenty-one of 36 males had a STAI-T score > 33.86 (population mean for males between ages 50-69 years) and nine of 24 females had a STAI-T score > 31.79 (population mean for females between ages 50-69 years).
- Fourteen had apathy scores ≥ 14.
- Forty-nine had minimal (0-13), seven had mild (14-19), three had moderate (20-28), and one had severe (29-63) symptoms of depression on BDI-II.
- Forty-four had an average PFS-16 score < 2.95 (i.e., fatigue not perceived as a significant problem).

Out of 48 subjects with PD included in the resting-state fMRI analysis:

- Six had a MoCA score < 26. None had a score < 21.
- Seventeen of 31 males had a STAI-T score > 33.86 (population mean for males between ages 50-69 years) and seven of 17 females had a STAI-T score > 31.79 (population mean for females between ages 50-69 years).
- Eleven had apathy scores ≥ 14.
- Forty-one had minimal (0-13), five had mild (14-19), one had moderate (20-28), and one had severe (29-63) symptoms of depression on BDI-II.
- 37 had an average PFS-16 score < 2.95 (i.e., fatigue not perceived as a significant problem).

**Correlations between the behavioral measures**

The STAI-T, BDI-II, apathy, and PFS-16 scores showed strong positive correlation with each other, but not with MDS-UPDRSIII. Tables 3a and 3b show the Spearman rho correlations and p values for 60 and 48 PD subjects, respectively.

**Table 3a. Correlations between behavioral measures (N=60)**

| | STAI-T | BDI-II | Apathy | PFS-16 | MDS-UPDRS III |
|---|---|---|---|---|---|
| STAI-T | 1 | | | | |
| BDI-II | 0.628 (p=0.000) | 1 | | | |
| Apathy | 0.524 (p=0.000) | 0.465 (p=0.000) | 1 | | |
| PFS-16 | 0.636 (p=0.000) | 0.721 (p=0.000) | 0.463 (p=0.000) | 1 | |
| MDS-UPDRS III | -0.071 (p=0.588) | -0.095 (p=0.472) | 0.119 (p=0.365) | 0.015 (p=0.908) | 1 |

**Table 3b. Correlations between behavioral measures (N=48)**

| | STAI-T | BDI-II | Apathy | PFS-16 | MDS-UPDRS III |
|---|---|---|---|---|---|
| STAI-T | 1 | | | | |
| BDI-II | 0.623 (p=0.000) | 1 | | | |
| Apathy | 0.532 (p=0.000) | 0.415 (p=0.003) | 1 | | |
| PFS-16 | 0.635 (p=0.000) | 0.664 (p=0.000) | 0.470 (p=0.001) | 1 | |
| MDS-UPDRS III | -0.034 (p=0.819) | -0.113 (p=0.446) | 0.012 (p=0.934) | 0.037 (p=0.803) | 1 |

**Table 4. Raw subcortical volumes**

| Volume ($mm^3$) | Con (N = 37) | PD (N = 60) |
|---|---|---|
| Total intracranial | 1,519,807 ± 170,334 | 1,588,293 ± 147,266 |
| Total brain | 1,126,181 ± 108,658 | 1,188,427 ± 107,975 |
| Gray matter | 612,695 ± 55,390 | 630,768 ± 55,336 |
| White matter | 449,229 ± 66,558 | 440,504 ± 150,971 |
| Cerebellum (GM) | | |
| Left | 52,293 ± 5,904 | 54,968 ± 5,913 |
| Right | 53,126 ± 5,839 | 57,471 ± 6,335 |
| Thalamus | | |
| Left | 6,945 ± 765 | 7,559 ± 859 |
| Right | 6,756 ± 824 | 7,237 ± 792 |
| Caudate | | |
| Left | 3,389 ± 400 | 3,366 ± 418 |

| | | | |
|---|---|---|---|
| | Right | 3,461 ± 423 | 3,465 ± 430 |
| Putamen | | | |
| | Left | 4,583 ± 722 | 4,405 ± 577 |
| | Right | 4,612 ± 562 | 4,559 ± 588 |
| Pallidum | | | |
| | Left | 2,007 ± 221 | 2,261 ± 322 |
| | Right | 1,920 ± 223 | 2,289 ± 335 |
| Accumbens | | | |
| | Left | 444 ± 98 | 324 ± 84 |
| | Right | 494 ± 87 | 452 ± 80 |
| Hippocampus | | | |
| | Left | 3,947 ± 355 | 3,907 ± 401 |
| | Right | 4,061 ± 400 | 4,119 ± 386 |
| Amygdala | | | |
| | Left | 1,608 ± 210 | 1,478 ± 250 |
| | Right | 1,729 ± 238 | 1,687 ± 196 |

GM: Gray matter

**Correlations between the behavioral data and subcortical volumes**

**Figure 1. Scatterplots**

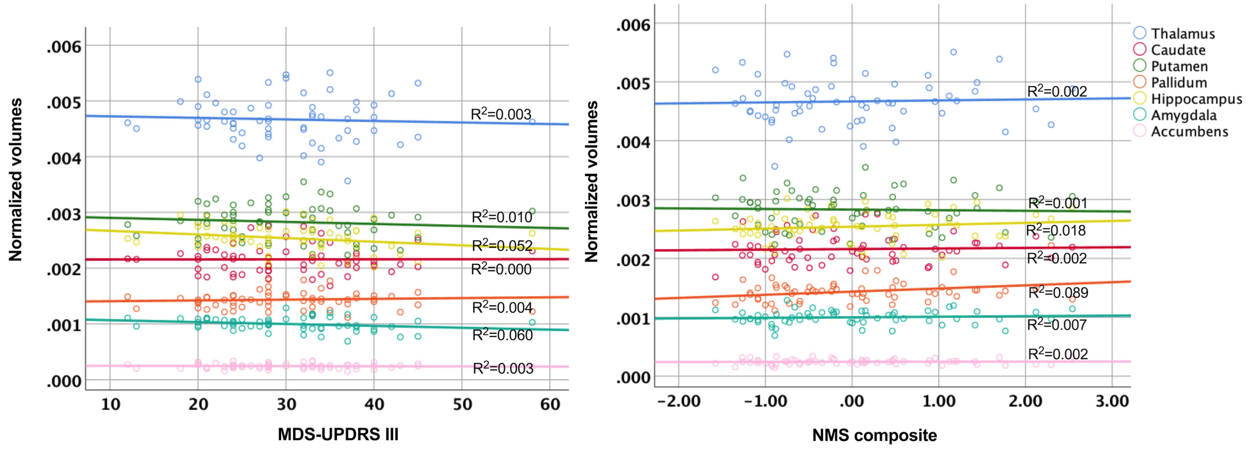


**Table 5. Amount of head motion during resting-state fMRI**

| | x (mean ± SD) | y (mean ± SD) | z (mean ± SD) |
|---|---|---|---|
| Mean translation (mm) | 0.23 ± 0.27 | 0.37 ± 0.54 | 0.26 ± 0.17 |
| Maximum translation (mm) | 0.48 ± 0.49 | 0.77 ± 0.85 | 0.65 ± 0.38 |
| Mean rotation (degree) | 0.38 ± 0.56 | 0.25 ± 0.27 | 0.22 ± 0.20 |
| Maximum rotation (degree) | 0.79 ± 0.94 | 0.53 ± 0.47 | 0.45 ± 0.36 |
| Scan-to-scan translation (mm) | 0.014 ± 0.007 | 0.071 ± 0.033 | 0.084 ± 0.056 |
| Scan-to-scan rotation (degree) | 0.034 ± 0.018 | 0.016 ± 0.008 | 0.015 ± 0.009 |

**Correlations between individual nonmotor symptom scores and pairwise functional connectivity**

See the interactive webpage https://bioimagesuiteweb.github.io/webapp/connviewer.html for the coordinates of the Shen Atlas nodes in the tables (Shen et al, 2013).

**APATHY**

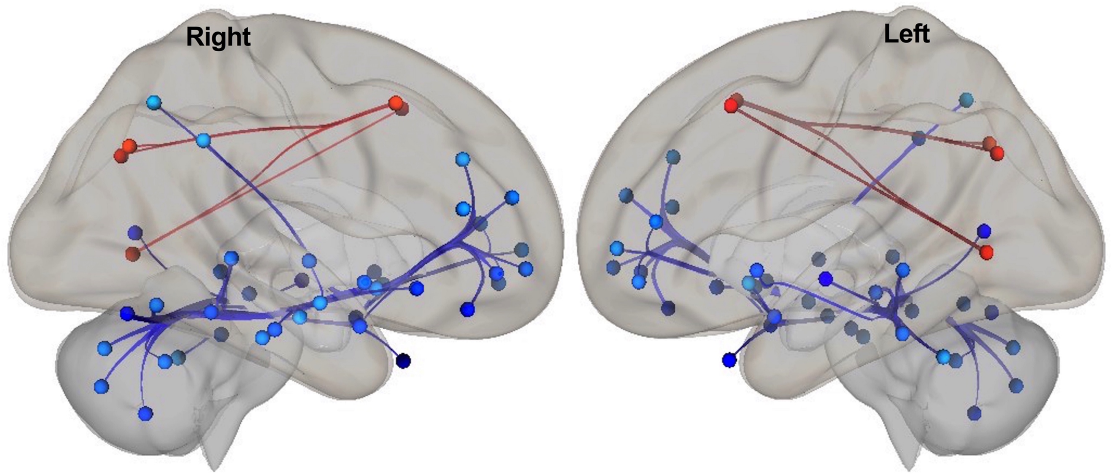


**Figure 2. Correlations between the apathy scores and pairwise functional connectivity** (FDR-corrected $p < 0.05$, warm colors: positive, cool colors: negative). Nodes in both hemispheres are displayed from right- and left-view on the MNI template.

**Table 6. Correlations between apathy scores and pairwise functional connectivity**

| **Negative Correlations** | | | | |
|---|---|---|---|---|
| *Node Pairs* | *Pair Labels* | | | |
| Numbers | **Node 1 (BA)** | **Node 2 (BA)** | T | p |
| (67)-(197) | R FG (BA37) | L STG (BA22) | -4.97 | 0.003 |
| (67)-(216) | R FG (BA37) | L V1 (BA17) | -4.19 | 0.017 |
| (67)-(93) | R FG (BA37) | R hippocampus | -4.00 | 0.021 |
| (125)-(11) | R putamen | R dlPFC (BA9) | -4.08 | 0.025 |
| (125)-(201) | R putamen | L FG (BA37) | -3.88 | 0.025 |
| (125)-(116) | R putamen | R cerebellum | -3.86 | 0.025 |
| (125)-(19) | R putamen | R dlPFC (BA46) | -3.84 | 0.025 |
| (125)-(8) | R putamen | R aPFC (BA10) | -3.76 | 0.025 |
| (125)-(113) | R putamen | R cerebellum | -3.71 | 0.025 |
| (201)-(36) | L FG (BA37) | R insula (BA13) | -4.16 | 0.026 |
| (201)-(34) | L FG (BA37) | R insula (BA13) | -3.98 | 0.026 |
| (201)-(169) | L FG (BA37) | L insula (BA13) | -3.84 | 0.026 |
| (201)-(64) | L FG (BA37) | R STG (BA22) | -3.75 | 0.027 |
| (67)-(233) | R FG (BA37) | L PHC (BA36) | -3.79 | 0.027 |
| (67)-(230) | R FG (BA37) | L hippocampus | -3.74 | 0.027 |
| (125)-(143) | R putamen | L aPFC (BA10) | -3.63 | 0.028 |
| (125)-(7) | R putamen | R aPFC (BA10) | -3.59 | 0.028 |
| (259)-(11) | L caudate | R dlPFC (BA9) | -4.04 | 0.028 |

| | | | | |
|---|---|---|---|---|
| (228)-(55) | L amygdala | R MTG (BA21) | -4.06 | 0.032 |
| (228)-(17) | L amygdala | R IFG (BA47) | -3.99 | 0.032 |
| (125)-(154) | R putamen | L dlPFC (BA46) | -3.44 | 0.035 |
| (125)-(238) | R putamen | L cerebellum | -3.41 | 0.035 |
| (125)-(47) | R putamen | R SMG (BA40) | -3.39 | 0.035 |
| (92)-(17) | R amygdala | R IFG (BA47) | -3.95 | 0.037 |
| (67)-(186) | R FG (BA37) | L TP (BA38) | -3.58 | 0.038 |
| (125)-(9) | R putamen | R aPFC BA10 | -3.33 | 0.039 |
| (93)-(94) | R hippocampus | R hippocampus | -3.81 | 0.039 |
| (93)-(66) | R hippocampus | R FG (BA37) | -3.74 | 0.039 |
| (93)-(59) | R hippocampus | R ITG (BA20) | -3.70 | 0.039 |
| (125)-(100) | R putamen | R cerebellum | -3.28 | 0.041 |
| (125)-(102) | R putamen | R cerebellum | -3.26 | 0.041 |
| (125)-(142) | R putamen | L aPFC (BA10) | -3.23 | 0.041 |
| (201)-(37) | L FG (BA37) | R insula BA13 | -3.52 | 0.045 |
| (125)-(43) | R putamen | R SPL (BA7) | -3.17 | 0.045 |
| (230)-(66) | L hippocampus | R FG (BA37) | -4.07 | 0.047 |
| (66)-(233) | R FG (BA37) | L PHC (BA36) | -3.74 | 0.047 |
| (228)-(70) | L amygdala | R FG (BA37) | -3.73 | 0.048 |
| ***Positive Correlations*** | | | | |
| *Node Pairs* | *Pair Labels* | | | |
| Numbers | **Node 1** | **Node 2** | T | p |
| (14)-(209) | R SFG (BA8) | L VA (BA19) | 4.33 | 0.022 |
| (209)-(149) | L VA (BA19) | L SFG (BA8) | 4.07 | 0.025 |
| (14)-(176) | R SFG (BA8) | L SPL (BA7) | 3.90 | 0.035 |
| (14)-(42) | R SFG (BA8) | R SPL (BA7) | 3.83 | 0.035 |

aPFC: Anterior prefrontal cortex, BA: Brodmann area, dlPFC: Dorsolateral prefrontal cortex, FG: Fusiform gyrus, IFG: Inferior frontal gyrus, orbital part, ITG: Inferior temporal gyrus, MTG: Middle temporal gyrus, PHC: Parahippocampus, SFG: Superior frontal gyrus, SMG: Supramarginal gyrus, SPL: Superior parietal lobule, STG: Superior temporal gyrus, TP: Temporal pole, V1: Primary visual area, V2: Secondary visual area, VA: Visual association area.
**Note**: Node 125 extends to the right nucleus accumbens.

**DEPRESSION**

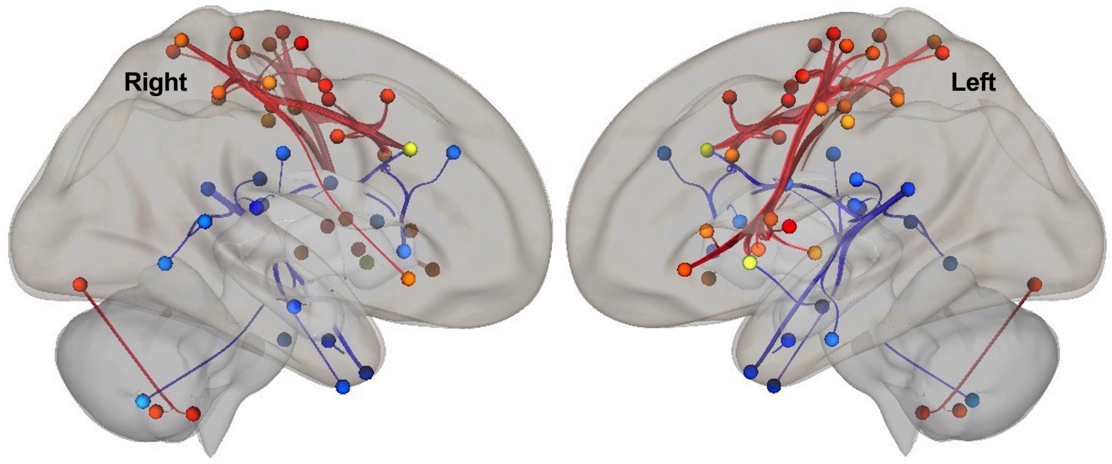

**Figure 3. Correlations between the BDI-II scores and pairwise functional connectivity** (FDR-corrected $p < 0.05$, warm colors: positive, cool colors: negative). Nodes in both hemispheres are displayed from right- and left-view on the MNI template.

**Table 7. Correlations between BDI-II scores and pairwise functional connectivity**

| ***Negative Correlations*** | | | | |
|---|---|---|---|---|
| *Node Pairs* | *Pair Labels* | | | |
| Numbers | **Node 1 (BA)** | **Node 2 (BA)** | T | p |
| (260)-(189) | L caudate | L TP (BA38) | -4.47 | 0.014 |
| (127)-(189) | R thalamus | L TP (BA38) | -4.18 | 0.018 |
| (123)-(15) | R caudate | R SFG (BA8) | -3.91 | 0.028 |
| (116)-(169) | R cerebellum | L insula (BA13) | -4.18 | 0.036 |
| (123)-(11) | R caudate | R dlPFC (BA9) | -4.17 | 0.036 |
| (123)-(221) | R caudate | L vACC (BA24) | -3.35 | 0.036 |
| (221)-(217) | L vACC (BA24) | L auditory | -3.29 | 0.040 |
| (264)-(189) | L thalamus | L TP (BA38) | -3.76 | 0.043 |
| (217)-(196) | L auditory | L ITG (BA20) | -4.03 | 0.044 |
| (217)-(69) | L auditory | R FG (BA37) | -3.86 | 0.045 |
| (217)-(235) | L auditory | L PHC (BA36) | -3.76 | 0.044 |
| (258)-(88) | L caudate | R dPCC (BA23) | -4.10 | 0.046 |
| (217)-(181) | L auditory | L SMG (BA40) | -3.64 | 0.047 |
| (217)-(15) | L auditory | R SFG (BA8) | -3.48 | 0.049 |
| (217)-(94) | L auditory | R hippocampus | -3.40 | 0.049 |
| (217)-(60) | L auditory | R ITG (BA20) | -3.39 | 0.049 |
| (217)-(65) | L auditory | R STG (BA22) | -3.39 | 0.049 |
| ***Positive Correlations*** | | | | |
| *Node Pairs* | *Pair Labels* | | | |
| Numbers | **Node 1 (BA)** | **Node 2 (BA)** | T | p |
| (38)-(221) | R S1 (BA1) | L vACC (BA24) | 5.28 | 0.001 |
| (38)-(155) | R S1 (BA1) | L dlPFC (BA46) | 5.04 | 0.001 |
| (38)-(15) | R S1 (BA1) | R SFG (BA8) | 4.78 | 0.002 |
| (221)-(26) | L vACC (BA24) | R PMC (BA6) | 4.40 | 0.009 |
| (151)-(41) | L IFG (BA47) | R SPL (BA7) | 4.35 | 0.012 |
| (151)-(175) | L IFG (BA47) | L SPL (BA7) | 4.31 | 0.012 |
| (38)-(28) | R S1 (BA1) | R SMA (BA6) | 4.00 | 0.016 |
| (109)-(76) | R cerebellum | R V2 (BA18) | 4.42 | 0.017 |
| (108)-(76) | R cerebellum | R V2 (BA18) | 4.18 | 0.018 |
| (221)-(39) | L vACC (BA24) | R S1 (BA1) | 3.89 | 0.021 |
| (221)-(32) | L vACC (BA24) | R PMC (BA6) | 3.85 | 0.021 |
| (221)-(166) | L vACC (BA24) | L PMC (BA6) | 3.83 | 0.021 |
| (15)-(41) | R SFG (BA8) | R SPL (BA7) | 4.10 | 0.023 |
| (221)-(160) | L vACC (BA24) | L PMC (BA6) | 3.68 | 0.025 |
| (221)-(33) | L vACC (BA24) | R S1 (BA1) | 3.67 | 0.025 |
| (221)-(179) | L vACC (BA24) | L SMG (BA40) | 3.61 | 0.026 |
| (41)-(155) | R SPL (BA7) | L dlPFC (BA46) | 3.91 | 0.027 |
| (221)-(167) | L vACC (BA24) | L S1 (BA1) | 3.55 | 0.028 |

| | | | | |
|---|---|---|---|---|
| (221)-(41) | L vACC (BA24) | R SPL (BA7) | 3.50 | 0.028 |
| (38)-(168) | R S1 (BA1) | L insula (BA13) | 3.70 | 0.030 |
| (38)-(170) | R S1 (BA1) | L insula (BA13) | 3.64 | 0.030 |
| (38)-(36) | R S1 (BA1) | R insula (BA13) | 3.60 | 0.030 |
| (221)-(172) | L vACC (BA24) | L S1 (BA1) | 3.44 | 0.030 |
| (38)-(169) | R S1 (BA1) | L insula (BA13) | 3.53 | 0.033 |
| (15)-(166) | R SFG (BA8) | L PMC (BA6) | 3.70 | 0.034 |
| (15)-(39) | R SFG (BA8) | R S1(BA1) | 3.67 | 0.034 |
| (38)-(84) | R S1(BA1) | R vACC (BA24) | 3.46 | 0.036 |
| (166)-(84) | L PMC (BA6) | R vACC (BA24) | 3.86 | 0.041 |
| (166)-(28) | L PMC (BA6) | R SMA (BA6) | 3.60 | 0.041 |
| (166)-(161) | L PMC (BA6) | L vACC (BA24) | 3.56 | 0.041 |
| (166)-(169) | L PMC (BA6) | L insula (BA13) | 3.55 | 0.041 |
| (166)-(168) | L PMC (BA6) | L insula (BA13) | 3.46 | 0.042 |
| (166)-(170) | L PMC (BA6) | L insula (BA13) | 3.44 | 0.042 |
| (261)-(166) | L putamen | L PMC (BA6) | 3.40 | 0.042 |
| (28)-(39) | R SMA (BA6) | R S1 (BA1) | 3.89 | 0.043 |
| (221)-(158) | L vACC (BA24) | L M1 (BA4) | 3.22 | 0.045 |
| (221)-(218) | L vACC (BA24) | L dPCC (BA31) | 3.20 | 0.045 |
| (151)-(179) | L IFG (BA47) | L SMG (BA40) | 3.68 | 0.047 |
| (151)-(171) | L IFG (BA47) | L S1 (BA1) | 3.64 | 0.047 |
| (160)-(163) | L PMC (BA6) | L PMC (BA6) | 3.87 | 0.049 |
| (160)-(169) | L PMC (BA6) | L insula (BA13) | 3.64 | 0.049 |
| (160)-(168) | L PMC (BA6) | L insula (BA13) | 3.62 | 0.049 |

**Note**: Node 217 is labeled as the left auditory node in Shen Atlas, however, it abuts the posterior horn of the lateral ventricle.

**FATIGUE**

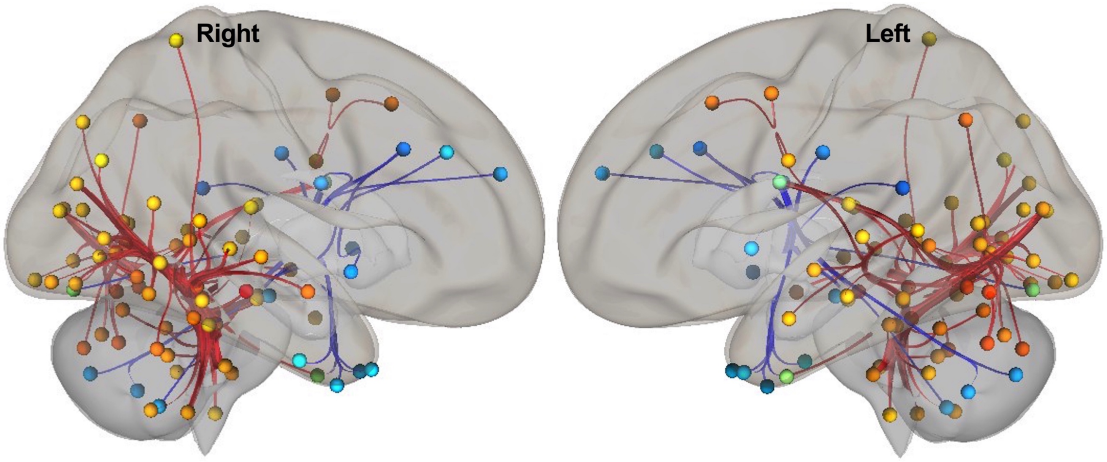


**Figure 4. Correlations between the PFS-16 scores and pairwise functional connectivity** (FDR-corrected $p < 0.05$, warm colors: positive, cool colors: negative). Nodes in both hemispheres are displayed from right- and left-view on the MNI template.

**Table 8. Correlations between PFS-16 scores and pairwise functional connectivity**

| ***Negative Correlations*** | | | | |
|---|---|---|---|---|
| *Node Pairs* | *Pair Labels* | | | |
| Numbers | **Node 1 (BA)** | **Node 2 (BA)** | T | p |
| (261)-(246) | L putamen | L cerebellum | -4.47 | 0.014 |
| (213)-(10) | L V2 (BA18) | R dlPFC (BA9) | -4.44 | 0.015 |
| (132)-(260) | R brainstem | L caudate | -3.63 | 0.016 |
| (260)-(51) | L caudate | R TP (BA38) | -3.88 | 0.026 |
| (260)-(60) | L caudate | R ITG (BA20) | -3.77 | 0.026 |
| (260)-(189) | L caudate | L TP (BA38) | -3.77 | 0.026 |
| (260)-(115) | L caudate | R cerebellum | -3.73 | 0.026 |
| (260)-(242) | L caudate | L cerebellum | -3.69 | 0.026 |
| (260)-(265) | L caudate | L brainstem | -3.56 | 0.026 |
| (260)-(202) | L caudate | L PHC (BA36) | -3.44 | 0.026 |
| (260)-(58) | L caudate | R ITG (BA20) | -3.39 | 0.026 |
| (15)-(34) | R SFG (BA8) | R insula (BA13) | -4.01 | 0.035 |
| (15)-(217) | R SFG (BA8) | L auditory | -3.96 | 0.035 |
| (103)-(11) | R cerebellum | R dlPFC (BA9) | -3.61 | 0.043 |
| (122)-(224) | R caudate | L dPCC (BA23) | -4.10 | 0.046 |
| ***Positive Correlations*** | | | | |
| *Node Pairs* | *Pair Labels* | | | |
| Numbers | **Node 1 (BA)** | **Node 2 (BA)** | T | p |
| 132)-(207) | R brainstem | L VA (BA19) | 5.01 | 0.002 |
| (132)-(72) | R brainstem | R VA (BA19) | 4.81 | 0.002 |
| (132)-(71) | R brainstem | R FG (BA37) | 4.78 | 0.002 |
| (132)-(209) | R brainstem | L VA (BA19) | 4.67 | 0.002 |
| (132)-(74) | R brainstem | R VA (BA19) | 4.42 | 0.003 |
| (132)-(68) | R brainstem | R FG (BA37) | 4.28 | 0.004 |
| (132)-(73) | R brainstem | R VA (BA19) | 4.22 | 0.004 |
| (132)-(205) | R brainstem | L VA (BA19) | 3.99 | 0.008 |
| (267)-(64) | L brainstem | R STG (BA22) | 4.43 | 0.009 |
| (267)-(63) | L brainstem | R STG (BA22) | 4.39 | 0.009 |
| (132)-(66) | R brainstem | R FG (BA37) | 3.86 | 0.011 |
| (132)-(210) | R brainstem | L V2 (BA18) | 3.80 | 0.012 |
| (113)-(177) | R cerebellum | L SPL (BA7) | 4.45 | 0.015 |
| (132)-(198) | R brainstem | L FG (BA37) | 3.67 | 0.015 |
| (132)-(98) | R brainstem | R V2 (BA18) | 3.61 | 0.016 |
| (127)-(73) | R thalamus | R VA (BA19) | 4.24 | 0.016 |
| (132)-(69) | R brainstem | R FG (BA37) | 3.58 | 0.016 |
| (132)-(206) | R brainstem | L VA (BA19) | 3.53 | 0.017 |
| (132)-(204) | R brainstem | L VA (BA19) | 3.50 | 0.017 |
| (132)-(192) | R brainstem | L MTG (BA21) | 3.49 | 0.017 |
| (237)-(204) | L cerebellum | L VA (BA19) | 4.34 | 0.018 |
| (103)-(204) | R cerebellum | L VA (BA19) | 4.15 | 0.018 |
| (251)-(204) | L cerebellum | L VA (BA19) | 4.05 | 0.018 |
| (133)-(204) | R brainstem | L VA (BA19) | 3.94 | 0.018 |

| | | | | |
|---|---|---|---|---|
| (267)-(204) | L brainstem | L VA (BA19) | 3.88 | 0.018 |
| (150)-(165) | L SFG (BA8) | L PMC (BA6) | 4.39 | 0.018 |
| (117)-(204) | R cerebellum | L VA (BA19) | 3.80 | 0.019 |
| (255)-(204) | L cerebellum | L VA (BA19) | 3.73 | 0.021 |
| (241)-(108) | L cerebellum | R cerebellum | 4.35 | 0.021 |
| (108)-(76) | R cerebellum | R V2 (BA18) | 3.98 | 0.024 |
| (108)-(213) | R cerebellum | R V2 (BA18) | 3.95 | 0.024 |
| (260)-(211) | L caudate | L V2 (BA18) | 4.05 | 0.026 |
| (260)-(191) | L caudate | L V2 (BA18) | 3.54 | 0.026 |
| (260)-(159) | L caudate | L PMC (BA6) | 3.50 | 0.026 |
| (260)-(216) | L caudate | L V1 (BA17) | 3.43 | 0.026 |
| (260)-(82) | L caudate | R V1 (BA17) | 3.42 | 0.026 |
| (260)-(215) | L caudate | L V1 (BA17) | 3.41 | 0.026 |
| (237)-(73) | L cerebellum | R VA(BA19) | 3.90 | 0.029 |
| (264)-(73) | L thalamus. | R VA(BA19) | 3.79 | 0.030 |
| (267)-(95) | L brainstem | R PHC (BA36) | 3.78 | 0.031 |
| (109)-(76) | R cerebellum | R V2 (BA18) | 4.08 | 0.033 |
| (108)-(76) | R cerebellum | R V2 (BA18) | 3.98 | 0.033 |
| (253)-(206) | L cerebellum | L VA (BA19) | 4.19 | 0.034 |
| (133)-(63) | R brainstem | R STG (BA22) | 3.99 | 0.036 |
| (133)-(204) | R brainstem | L VA (BA19) | 3.94 | 0.036 |
| (133)-(74) | R brainstem | R VA (BA19) | 3.77 | 0.036 |
| (133)-(54) | R brainstem | R MTG (BA21) | 3.62 | 0.036 |
| (133)-(64) | R brainstem | R STG (BA22) | 3.60 | 0.036 |
| (133)-(65) | R brainstem | R STG (BA22) | 3.59 | 0.036 |
| (108)-(213) | R cerebellum | L V2 (BA18) | 3.95 | 0.037 |
| (267)-(54) | L brainstem | R MTG (BA21) | 3.61 | 0.040 |
| (267)-(74) | L brainstem | R VA (BA19) | 3.56 | 0.040 |
| (108)-(80) | R cerebellum | R V2 (BA18) | 3.64 | 0.042 |
| (108)-(81) | R cerebellum | R V2 (BA18) | 3.60 | 0.042 |
| (253)-(232) | L cerebellum | L hippocampus | 3.90 | 0.043 |
| (103)-(72) | R cerebellum | R VA (BA19) | 3.70 | 0.043 |
| (103)-(210) | R cerebellum | L V2 (BA18) | 3.66 | 0.043 |
| (103)-(73) | R cerebellum | R VA(BA19) | 3.59 | 0.043 |
| (133)-(50) | R brainstem | R AG (BA39) | 3.48 | 0.043 |
| (132)-(41) | R brainstem | R SPL (BA7) | 3.14 | 0.044 |
| (132)-(75) | R brainstem | R VA (BA19) | 3.10 | 0.044 |
| (132)-(49) | R brainstem | R AG (BA39) | 3.10 | 0.044 |
| (247)-(66) | L cerebellum | R FG (BA37) | 3.99 | 0.046 |
| (66)-(202) | R FG (BA37) | L PHC (BA36) | 3.74 | 0.046 |
| (267)-(190) | L brainstem | L MTG (BA21) | 3.44 | 0.049 |
| (132)-(197) | R brainstem | L STG (BA22) | 3.04 | 0.049 |
| (132)-(76) | R brainstem | R V2 (BA18) | 3.03 | 0.049 |
| (132)-(191) | R brainstem | L MTG (BA21) | 3.01 | 0.049 |
| (260)-(192) | L caudate | L MTG (BA21) | 3.13 | 0.049 |
| (260)-(79) | L caudate | R V2 (BA18) | 3.12 | 0.049 |

| | | | | |
|---|---|---|---|---|
| (108)-(70) | R cerebellum | R FG (BA37) | 3.45 | 0.049 |
| (108)-(214) | R cerebellum | L V2 (BA18) | 3.43 | 0.049 |
| (266)-(204) | L brainstem | L VA (BA19) | 3.33 | 0.049 |
| (248)-(204) | L cerebellum | L VA (BA19) | 3.31 | 0.049 |

AG: Angular gyrus, BA: Brodmann area, dlPFC: Dorsolateral prefrontal cortex, dPCC: Dorsal posterior cingulate cortex, FG: Fusiform gyrus, ITG: Inferior temporal gyrus, MTG: Middle temporal gyrus, PHC: Parahippocampus, PMC: Premotor cortex, SFG: Superior frontal gyrus, SMG: Supramarginal gyrus, SPL: Superior parietal lobule, STG: Superior temporal gyrus, TP: Temporal pole, V1: Primary visual area, V2: Secondary visual area, VA: Visual association area.
**Note**: Nodes 132, and 133/267 correspond to mid brain and pons, respectively. Node 217 is labeled as the left auditory node in Shen Atlas, however, it abuts the posterior horn of the lateral ventricle.

**ANXIETY**

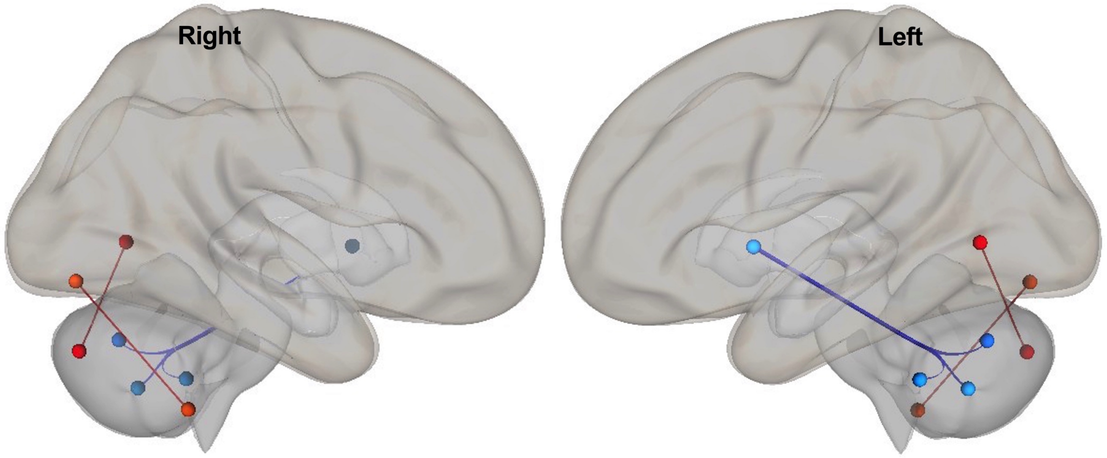


**Figure 5. Correlations between the STAI-T scores and pairwise functional connectivity** (FDR-corrected $p < 0.05$, warm colors: positive, cool colors: negative). Nodes in both hemispheres are displayed from right- and left-view on the MNI template.

**Table 9. Correlations between STAI-T scores and pairwise functional connectivity**

| ***Negative Correlations*** | | | | |
|---|---|---|---|---|
| *Node Pairs* | *Pair Labels* | | | |
| Numbers | **Node 1 (BA)** | **Node 2 (BA)** | T | p |
| (261)-(252) | L putamen | L cerebellum | -4.14 | 0.035 |
| (261)-(246) | L putamen | L cerebellum | -3.86 | 0.035 |
| (261)-(253) | L putamen | L cerebellum | -3.83 | 0.035 |
| ***Positive Correlations*** | | | | |
| *Node Pairs* | *Pair Labels* | | | |
| Numbers | **Node 1 (BA)** | **Node 2 (BA)** | T | p |
| (108)-(76) | R cerebellum | R V2 (BA18) | 4.54 | 0.011 |
| (111)-(209) | R cerebellum | L VA (BA19) | 4.31 | 0.024 |

BA: Brodmann area, V2: Secondary visual area, VA: Visual association area.
**Note:** Node 261 extends to the left globus pallidus.
